# Predicting sediment and nutrient concentrations from high-frequency water-quality data

**DOI:** 10.1101/599712

**Authors:** Catherine Leigh, Sevvandi Kandanaarachchi, James M. McGree, Rob J. Hyndman, Omar Alsibai, Kerrie Mengersen, Erin E. Peterson

## Abstract

Water-quality monitoring in rivers often focuses on the concentrations of sediments and nutrients, constituents that can smother biota and cause eutrophication. However, the physical and economic constraints of manual sampling prohibit data collection at the frequency required to adequately capture the variation in concentrations through time. Here, we developed models to predict total suspended solids (TSS) and oxidized nitrogen (NOx) concentrations based on high-frequency time series of turbidity, conductivity and river level data from *in situ* sensors in rivers flowing into the Great Barrier Reef lagoon. We fit generalized-linear mixed-effects models with continuous first-order autoregressive correlation structures to water-quality data collected by manual sampling at two freshwater sites and one estuarine site and used the fitted models to predict TSS and NOx from the *in situ* sensor data. These models described the temporal autocorrelation in the data and handled observations collected at irregular frequencies, characteristics typical of water-quality monitoring data. Turbidity proved a useful and generalizable surrogate of TSS, with high predictive ability in the estuarine and fresh water sites. Turbidity, conductivity and river level served as combined surrogates of NOx. However, the relationship between NOx and the covariates was more complex than that between TSS and turbidity, and consequently the ability to predict NOx was lower and less generalizable across sites than for TSS. Furthermore, prediction intervals tended to increase during events, for both TSS and NOx models, highlighting the need to include measures of uncertainty routinely in water-quality reporting. Our study also highlights that surrogate-based models used to predict sediments and nutrients need to better incorporate temporal components if variance estimates are to be unbiased and model inference meaningful. The transferability of models across sites, and potentially regions, will become increasingly important as organizations move to automated sensing for water-quality monitoring throughout catchments.

## Introduction

Measuring the concentrations of sediments and nutrients in rivers, and understanding how they change through time, is a major focus of water-quality monitoring given the potential detrimental effects these constituents have on aquatic ecosystems. Such knowledge can help inform the effective management of our land, waterways and oceans, including World Heritage Areas such as the Great Barrier Reef in the Australian tropics [1,2,3]. In regions dominated by highly seasonal, event-driven climates, such as those in the tropics, high-magnitude wet-season flows can transport large quantities of sediments and nutrients from the land downstream in relatively short time frames [4]. The rapidity of change in sediment and nutrient concentrations during high-flow events poses challenges for water-quality monitoring based on discrete manual sampling of water followed by laboratory measurement of concentrations, which is time consuming, costly and typically temporally sparse. Relatively low sampling frequency increases the chances of missing water-quality events, but high flows may preclude the safety conditions required for manual sampling, and sample collection at the frequency required to capture change in concentrations may not always be physically or economically practical. The spatial sparsity of measurements from manual sampling is also problematic. For example, the Great Barrier Reef lagoon stretches over 3000 km of coastline, but the data currently used to validate estimates of sediments and nutrients flowing to the lagoon are collected from just 43 sites [5]. This lack of data limits knowledge and understanding of sediments and nutrient concentrations in both space and time.

*In situ* sensors have the potential to complement or circumvent the need for manual sampling and laboratory analysis, whilst also providing monitoring data at the frequencies required to capture the full range of water-quality conditions occurring in rivers (e.g. every 15-60 mins). However, *in situ* sensors currently used to measure sediments and/or nutrients (e.g. nitrate) have drawbacks related to biofouling and drift, excessive power requirements, high costs, and/or low accuracy and precision [6]. An alternative is to use *in situ* sensors to measure water-quality variables, such as turbidity, conductivity, and river level (i.e. height), that have the potential to act individually or in combination as surrogates for sediments and nutrients [7,8]. However, an in-depth understanding of the relationship between sediment and nutrient dynamics and other water-quality variables is needed before the latter can be used as surrogate measures.

Turbidity is a visual property of water indicative of its clarity (or lack thereof) due to suspended particles of abiotic and biotic origin that absorb and scatter light. As a result, turbidity tends to increase during high-flow events in rivers, when waters often contain high concentrations of particles (e.g. sediments and nutrients from runoff-derived soil erosion), which makes it a popular surrogate for total suspended solids (TSS; e.g. [4]). Turbidity can also increase when water residence times increase during low flows, due to the resultant concentration of suspended particles, or when high concentrations of microalgae reduce water clarity. Both turbidity and conductivity of water can change rapidly during flow events. Conductivity reflects the ability of water to pass an electric current as determined by the concentration of ions including nitrate and nitrite (oxidized nitrogen; NOx= nitrite + nitrate). As such, new inputs of fresh water will typically decrease conductivity in rivers as waters rapidly dilute. In contrast, conductivity tends to increase during low-flow periods and when water levels decline. Turbidity and conductivity, together with river level, thus have the potential to act as a combined proxy for nutrients such as NOx (e.g. [9]).

A wide range of modelling techniques have been used to describe the relationship between sediment and nutrient concentrations and other water quality constituents. For example, Artificial Neural Networks have been used to predict nitrate from multiple measures including other nutrient concentrations [10], and standard major axis regression to quantify the relationship between turbidity and TSS [11]. One of the most common approaches is to use linear regression to predict TSS from turbidity [4,7,8,12,13,14,15], nitrogen species from turbidity and conductivity [4,9] and phosphorus species from turbidity [4,7,8,12,13,14,16]. However, these regression models typically fail to account for the temporal autocorrelation (i.e. serial correlation) inherent in water-quality time series and/or the heteroscedasticity in the data. This violates the underlying assumptions (i.e. identical and independently distributed residuals), which can lead to biased variance estimates, inflated statistical significance of predictor variables and thus incorrect inference. Models that account for temporal autocorrelation and heteroscedasticity through the incorporation of random effects and/or specific variance-covariance structures can produce more accurate and precise predictions when temporal correlation exists in the data (e.g. [17,18]). Despite the advantages of using mixed-effects models that account for temporal correlation to predict concentrations of sediments and nutrients from water-quality time series, they are rarely used for this purpose.

Our key objective was to predict TSS and NOx from high-frequency water-quality data using models that accounted explicitly for temporal autocorrelation and heteroscedasticity. We used turbidity (NTU), conductivity (µS/cm) and river level (m) data collected using *in situ* sensors in rivers flowing into the Great Barrier Reef lagoon (Fig 1), along with water-quality data measured in the laboratory, as surrogate covariates. We aimed to assess whether relationships between TSS or NOx and the water-quality surrogates differed (i) among sites and (ii) between estuarine and fresh waters. Surrogate approaches are often site-specific and as such suffer from lack of transferability [15]. Thus, we further aimed to assess (iii) whether a single mixed-effects model fit to the water quality surrogates could be used to predict TSS or NOx over multiple locations, and when using data collected by *in situ* sensors. By investigating the predictive ability of the models, our findings will provide a basis to determine the most effective water-quality surrogates for TSS and NOx, along with the potential generalizability of the models across locations in the study area.

**Fig 1.**
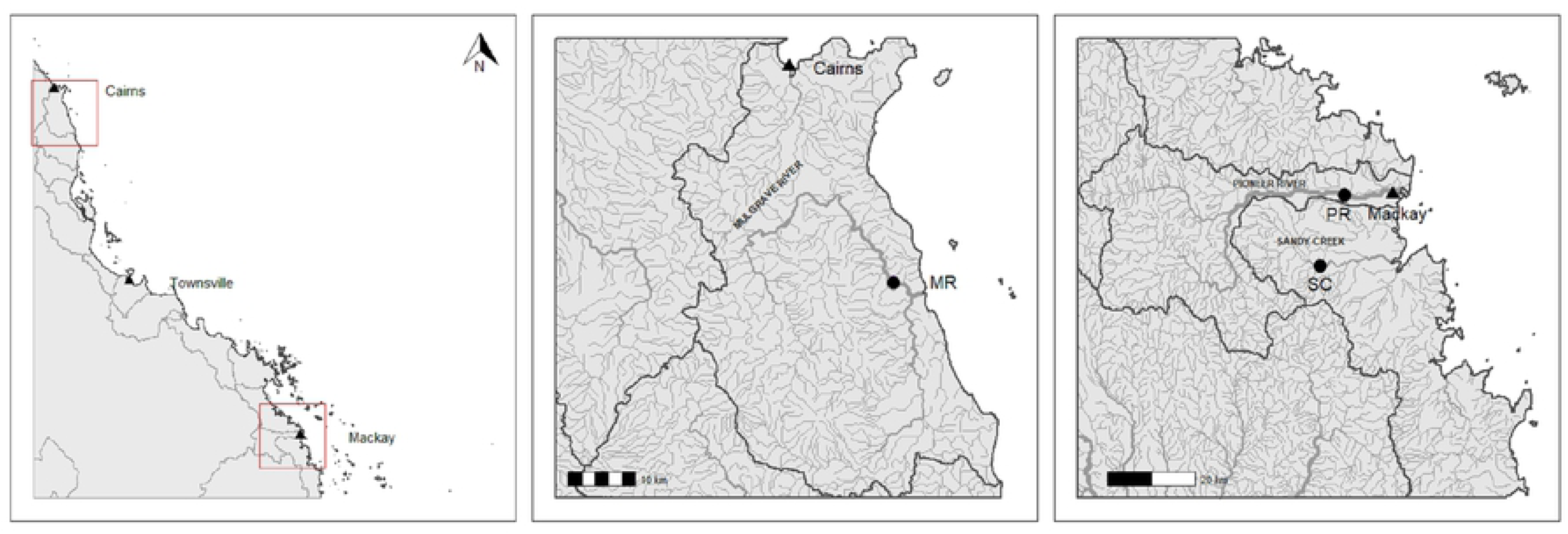
The study region. Study sites (closed circles), rivers and catchment boundaries within north tropical Queensland, Australia (left panel), the Wet Tropics (MR; middle panel) and Mackay Whitsunday regions (PR and SC; right panel). Closed triangles show the major towns of Cairns, Townsville and Mackay.

## Materials and methods

### Study region and sites

Our three study sites are located in rivers that flow into the Great Barrier Reef lagoon along the northeast coast of tropical Australia in Queensland (Fig 1). Two of the sites (Sandy Creek and Pioneer River) lie within the Mackay Whitsunday region and the third (Mulgrave River) lies within the Wet Tropics region. These two regions are characterized by seasonal climate, with higher rainfall and air temperatures in the ‘wet’ season and lower rainfall and air temperatures in the ‘dry’ season. Although there is inter-annual seasonal variation in climate and river flow in both regions, the wet season typically occurs from December to April in the Mackay Whitsunday region, and from November to April in the more northern Wet Tropics region [19,20,21]. The wet season is typically associated with tropical cyclones, monsoonal rainfall and associated event-flows in rivers, and the dry season with low to zero surface flow.

Pioneer River rises in the forested uplands of the Great Dividing Range in north Queensland [19]. Many of its upper reaches lie within National or State Parks, whilst land use in the mid and lower reaches is dominated by sugarcane farming. Sandy Creek is a low-lying coastal-plain stream south of the Pioneer River, where the dominant land use is also sugarcane farming. The Mulgrave River in the Wet Tropics World Heritage Area rises, like the Pioneer River, in forested National Park uplands of the Great Dividing Range and flows through mostly cleared alluvial floodplains in its lower reaches [22]. The Pioneer River and Sandy Creek sites (PR and SC) are in freshwater reaches, and the Mulgrave River site (MR) is in an estuarine reach. The monitored catchment area of each site is 1466 km^2^ (PR), 326 km^2^ (SC) and 789 km^2^ (MR). We chose these sites because they had comprehensive water-quality datasets available containing both laboratory-measured sediment and nutrient concentrations, as well as high frequency *in situ* water-quality data from multiple sensors.

### Laboratory and *in situ* sensor data

The Queensland Department of Environment and Science (DES) has installed an *in situ* automated water-quality sensor (YSI EXO2 Sonde attached with an EXO Turbidity Smart Sensor 599101-01 and EXO Conductivity & Temperature Smart Sensor 599870) at each of the three study sites. Sensors are housed in a flow cell in water-quality monitoring stations on riverbanks; water is pumped at regular intervals from the river to the flow cell, every hour or hour and a half depending on the site and variable being measured, and sometimes more frequently during event flows. The sensors measure and record turbidity (NTU) and electrical conductivity at 25 °C (conductivity; µS/cm). Pressure-induction sensors record river level (i.e. height in meters from the riverbed to the water surface; level, m) every 10 minutes. Time-matched observations of level for the occasional out-of-step turbidity or conductivity measurement are provided via linear interpolation of the ten-minute level data. All data were quality controlled and assured, including detection and removal of technical anomalies [23], prior to analysis.

DES manually collect grab-samples of water approximately monthly, and more frequently during event flows in the wet season when safety permits, from each site for laboratory analysis of water quality, as part of their Great Barrier Reef Catchment Loads Monitoring Program [24]. The laboratory data therefore contain unequally spaced observations of water quality through time. The goal of the program is to track long-term trends in the quality of water entering the Great Barrier Reef lagoon from adjacent catchments, as part of the Paddock to Reef program [25]. Collection, storage and transport of grab-samples is conducted under strict quality control and assurance procedures [26,27,28]. Samples are analyzed in the National Association of Testing Authorities credited Science Division Chemistry Centre laboratories (Dutton Park, Queensland) for turbidity, conductivity and concentrations of TSS (mg/L) and NOx (mg/L) following standard methods [29]. DES also record river level on-site on most occasions when grab-samples are collected.

Turbidity, conductivity, TSS and NOx data measured in the laboratory were available from January 2016 to June 2017 at SC and MR and from January 2016 to October 2017 at PR (Table 1, Figs 2–4). Turbidity, conductivity and level data measured and recorded *in situ* by automated sensors were available from March 2017 to March 2018 at all three sites (Table 1, Figs 2–4). Inputs of saline water from groundwater inputs or tidal influence will increase the conductivity of surface waters and confound the relationship between conductivity and NOx. For this reason, all conductivity observations under tidal influence at MR (i.e. those greater than the maximum conductivity observed across the two freshwater sites over the same time span, which was 1100 µS/cm) were removed from the laboratory and sensor data prior to analysis. The ranges of the laboratory-measured turbidity, conductivity and level data at each site were comparable and observations exhibited similar patterns through time as the respective turbidity, conductivity and level data from the *in situ* sensors at each site (Figs 2–4, Table 1), validating the use of the laboratory data to build models for subsequent prediction using the sensor data.

**Table 1.**
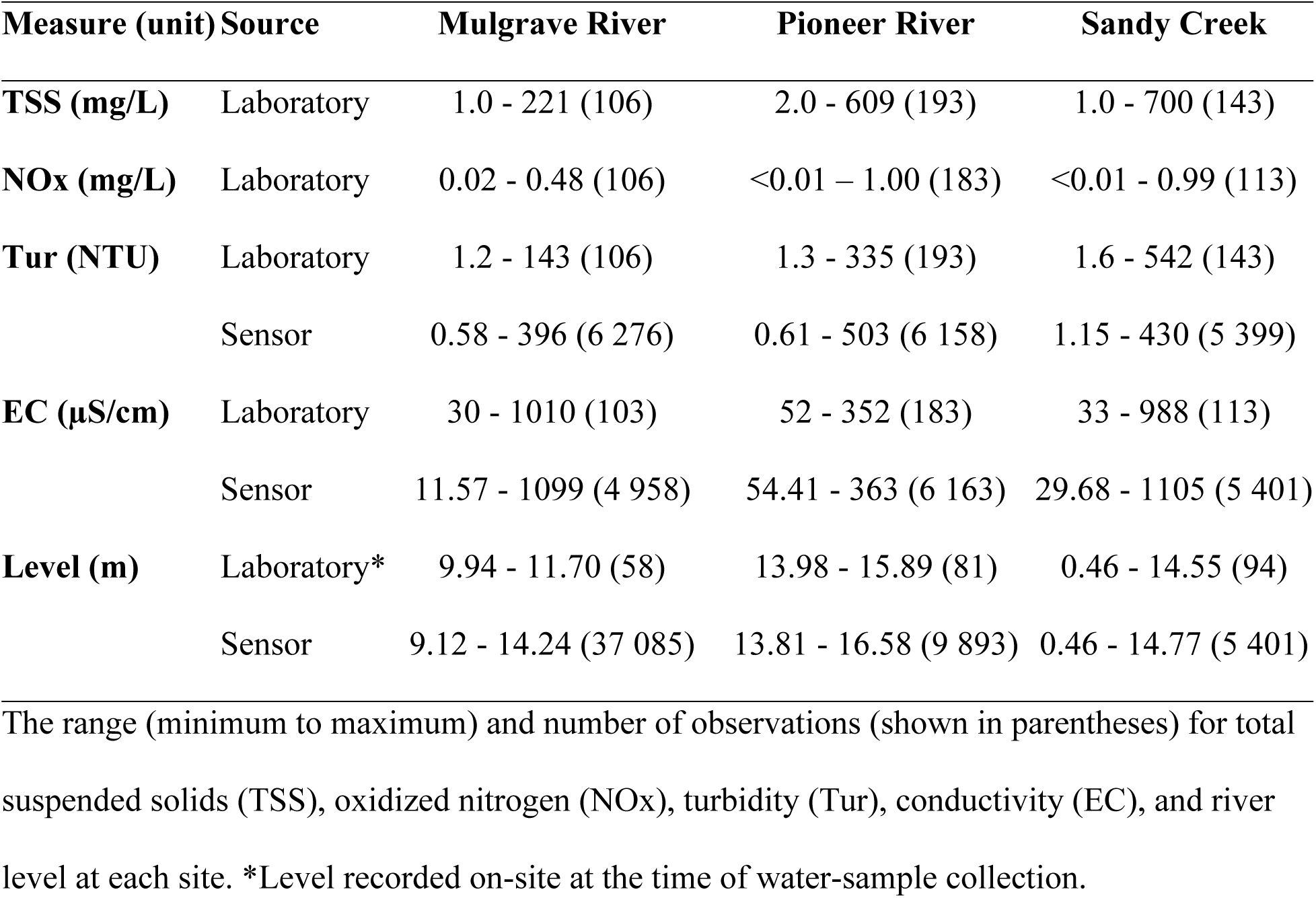
Water quality at each study site.

**Fig 2.**
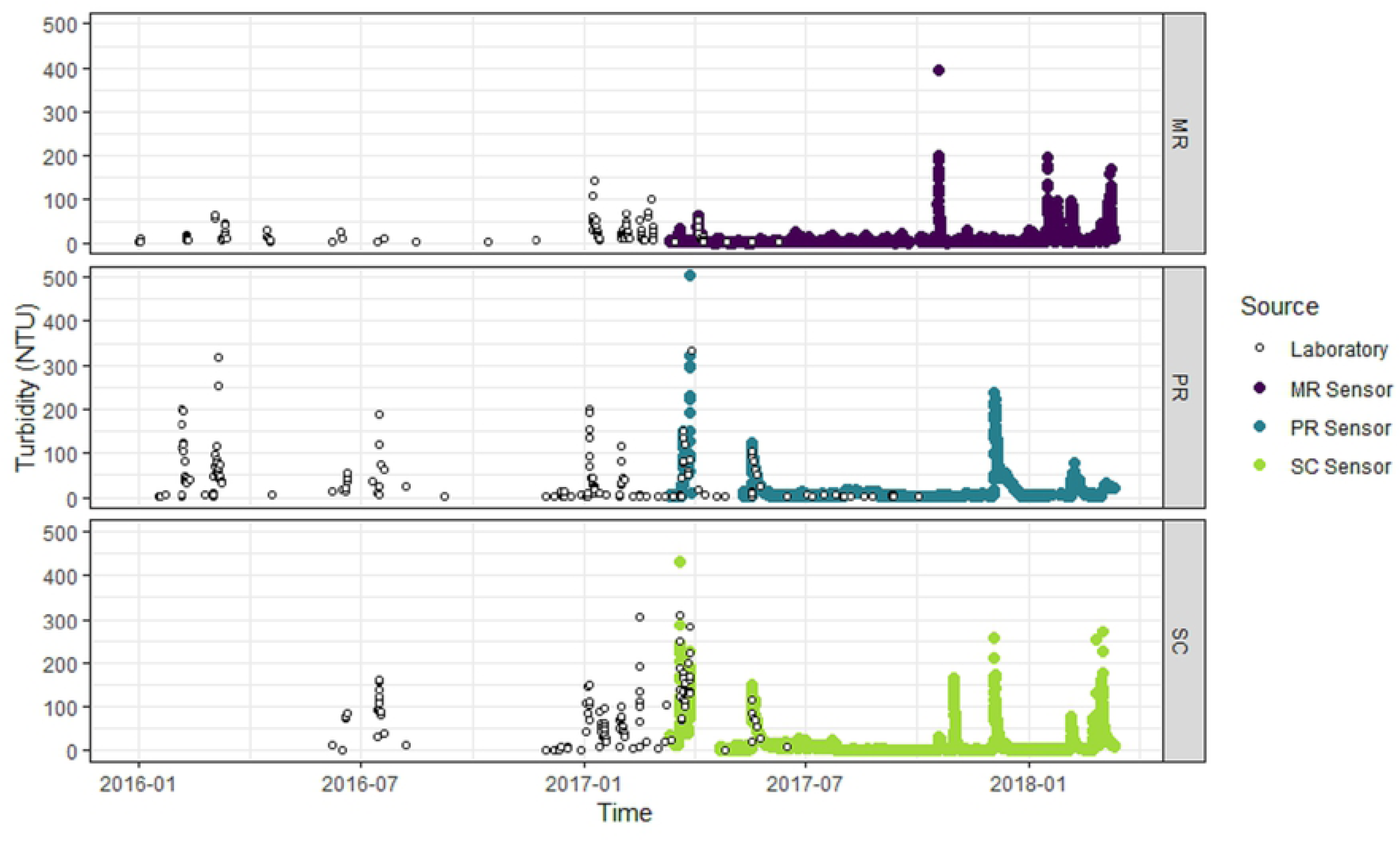
Turbidity of water at each study site. Laboratory-measured (open circles) and *in situ* sensor-measured turbidity (NTU). Mulgrave River (MR; purple points), Pioneer River (PR, blue points) and Sandy Creek (SC; light green points).

**Fig 3.**
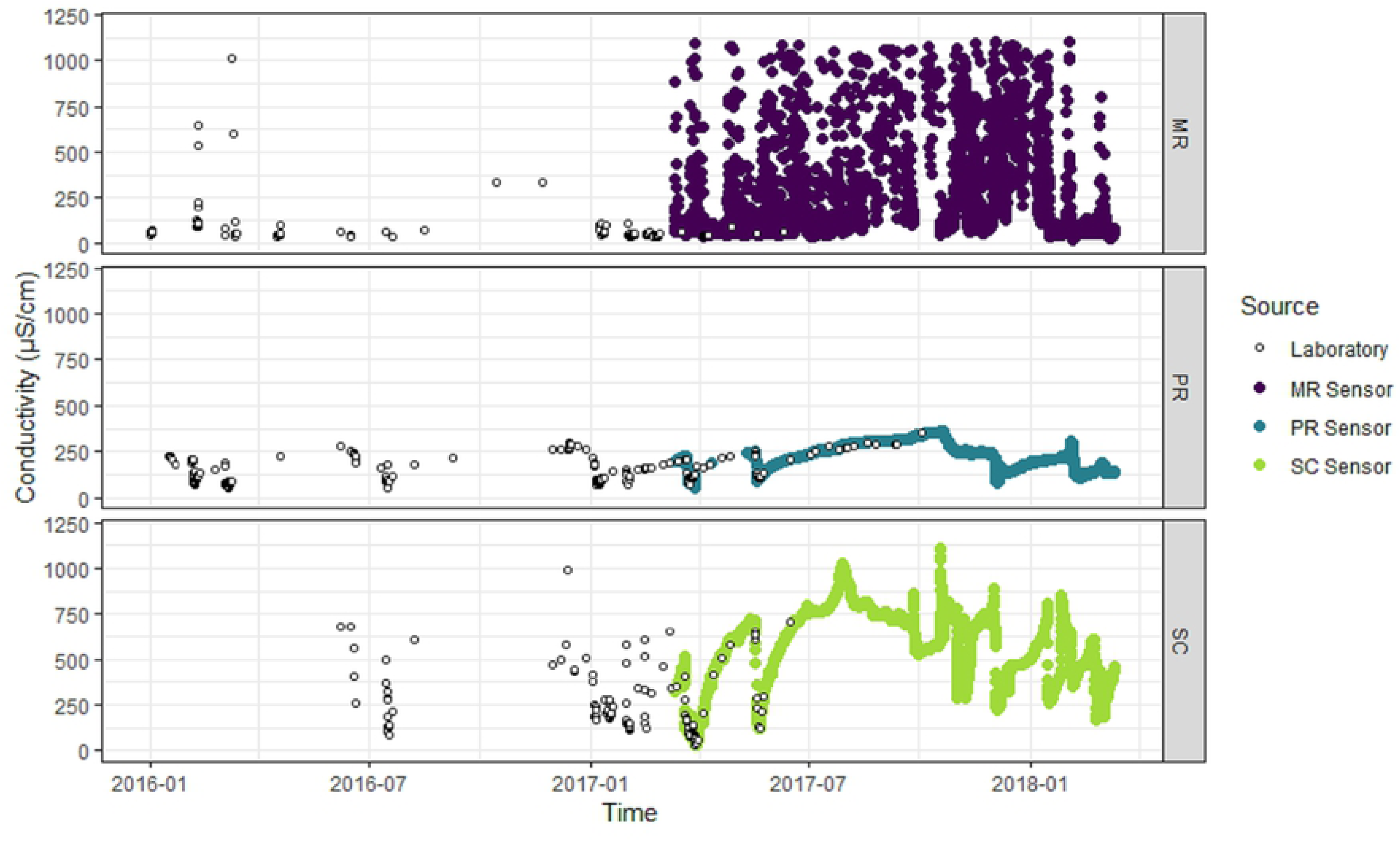
Conductivity of water at each study site. Laboratory-measured (open circles) and *in situ* sensor-measured conductivity (µS/cm) at Mulgrave River (MR; purple points), Pioneer River (PR, blue points) and Sandy Creek (SC; light green points).

**Fig 4.**
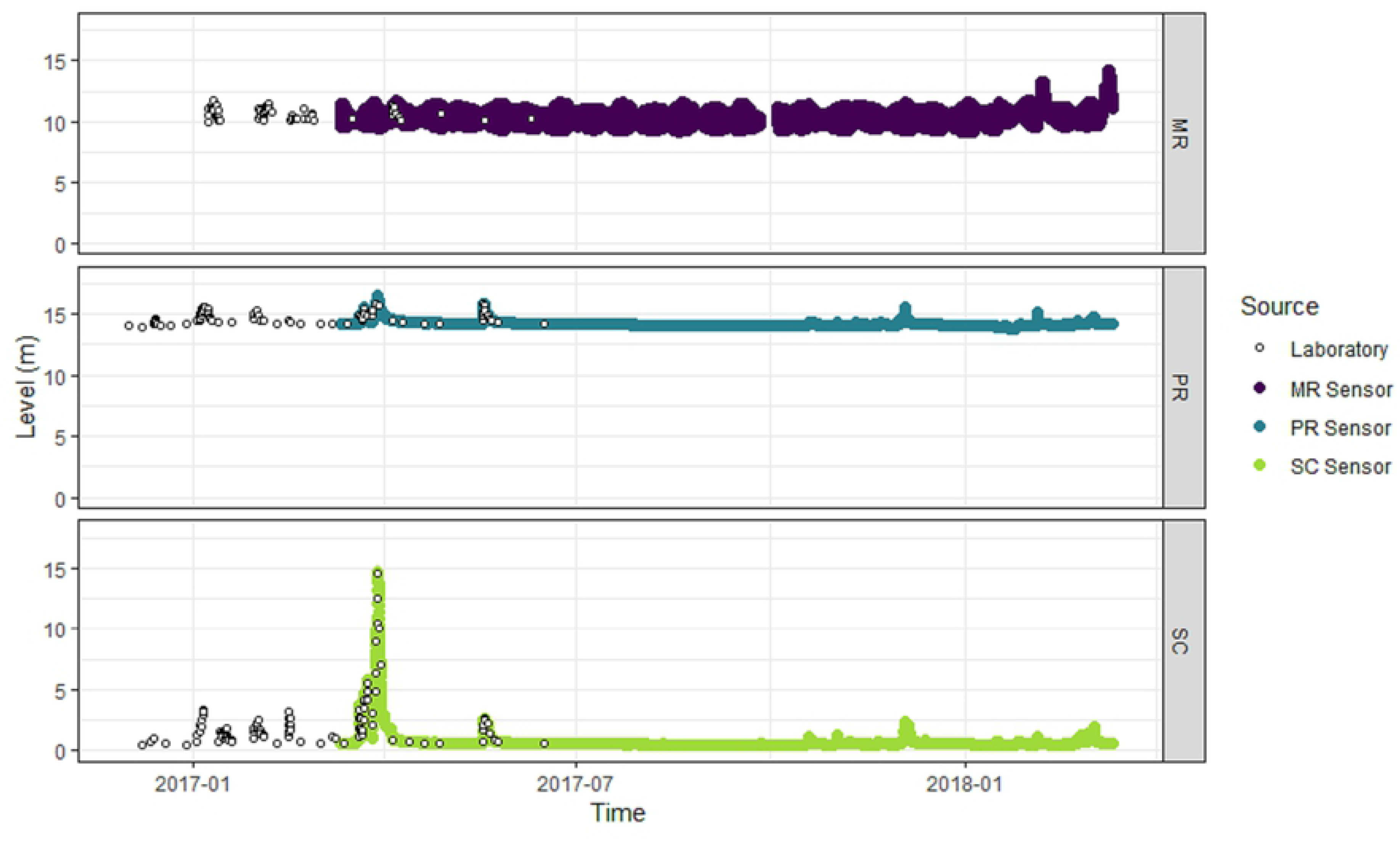
Height of water at each study site. River level (m) measured on-site at the time of water sample collection (open circles) and by *in situ* sensors at Mulgrave River (MR; purple points), Pioneer River (PR, blue points) and Sandy Creek (SC; light green points).

### Statistical analysis

We fit generalized-linear mixed-effects models with a continuous first-order autoregressive correlation (AR(1)) structure [30] to the laboratory TSS or NOx (i.e. the response) and surrogate water-quality variables (i.e. the covariates). The models are of the form:

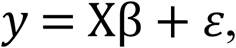

where *y* is an *n*-dimensional vector of TSS or NOx collected at times *t*_1_,…,*t*_*n*_; *n* is the number of observations; X is an *n* × *p* design matrix of *p* covariates collected at times *t*_1_,…,*t*_*n*_;β is a *p*-dimensional vector of estimated regression coefficients; and *ε* is an *n*-dimensional vector of zero-mean, normally distributed errors with covariance matrix *σ*^2^Λ. The covariance is defined by a continuous AR(1) structure, such that *Corr*(*ε*_*t*_, *ε*_*t* − 1_) = *ϕ*^*t_i_* − *t*_*i* − 1_^, where *ϕ* is the parameter in the AR(1) process, which can range between 0 and 1 and defines how the autocorrelation declines with time. The continuous AR(1) structure accounts for both the temporal correlation and unequal temporal spacing present in the laboratory time-series data.

We selected potential covariates for the TSS and NOx models based on plausible mechanisms that could cause changes in TSS or NOx, evidence from the literature, exploratory data analysis, and the availability of covariates within the laboratory dataset (Fig 5, Step 1; [31]).

**Fig 5:**
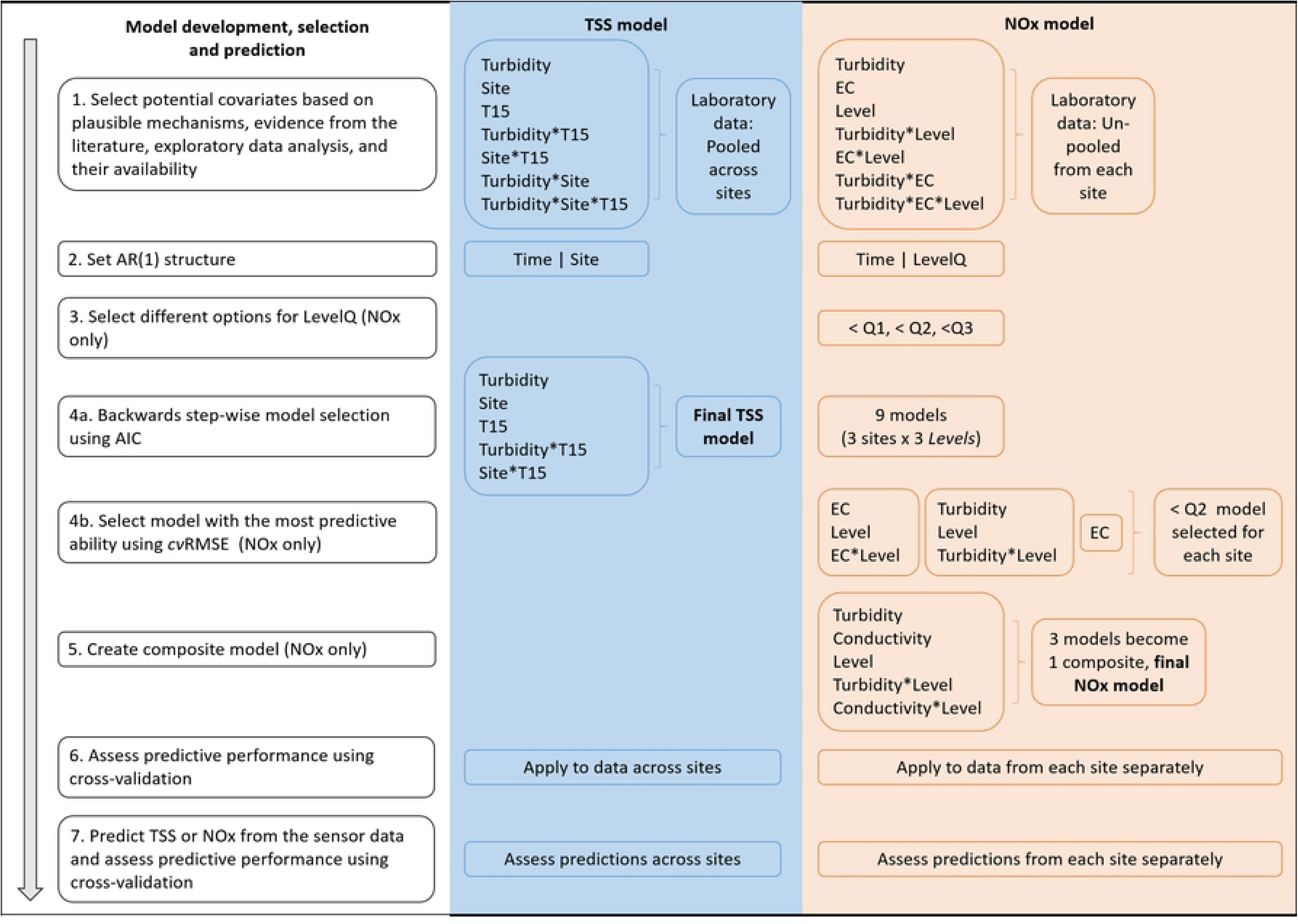
Model development, selection and prediction for the final total suspended solids (TSS) and oxidized nitrogen (NOx) models. LevelQ is a categorical variable with two levels based on first, second or third quartiles of the data (Q1, Q2 or Q3). Turbidity, conductivity and level covariates were all log_10_-transformed prior to analysis.

#### TSS model

Exploratory analyses indicated that including data from all three sites in a single TSS model was appropriate; there was a strong and similar positive relationship between turbidity and TSS at all sites (S1 Figure), reflecting the physical properties of these variables and the processes underlying water quality dynamics in rivers [32,33]. The suite of covariates we selected for the TSS model included: turbidity measured in the laboratory; T15, a categorical variable representing turbidity < 15 NTU (‘below’) or ≥ 15 NTU (‘above’); site (MR, PR, SC); and all of their interactions (Fig 5, Step 1). We also included site as a grouping variable in the temporal correlation structure, to account for within-site correlation (Fig 5, Step 2). We included T15 because the intercept for the relationship with TSS appeared to change below 15 NTU, particularly at freshwater sites PR and SC (S1 Figure). In addition, 15 NTU is the water-quality guideline value for turbidity in freshwater streams and rivers in northern Australia [34].

We used a two-step model-selection process to identify the final TSS model. First, we implemented a backwards-stepwise model-selection procedure to identify the subset of covariates that had the most support in the data (Fig 5, Step 4a). Parameters were estimated using maximum likelihood and models were compared using the Akaike Information Criterion (AIC) [35]. Next, we assessed the predictive performance of the model using a 5-fold cross-validation (*cv*) procedure designed specifically for temporally correlated data (Fig 5, Step 6) [36]. Validation data from each site were created by dividing the time series into five blocks of chronologically ordered observations. Maximum likelihood can produce biased estimates of *ϕ* [37] and so the final model was iteratively refit without the validation data, using restricted maximum likelihood (REML) for parameter estimation. We then used the observations from the validation set and the associated cross-validation predictions to calculate the root mean-square error (cvRMSE) and the 95% prediction coverage value (cvPC). An r-squared statistic (*cvR^2^*) was also generated as the squared Spearman rank correlation between the observations and the cross-validated predictions.

#### NOx model

Relationships between NOx and the potential covariates conductivity, turbidity (both measured in the laboratory) and river level (measured on-site at the time of water sampling) were more complex and site-specific than the relationship between turbidity and TSS (S2-S4 Figures). Therefore, we first fit NOx models for each site separately (Fig 5, Step 1). The models included covariates for conductivity, turbidity, level and all of their interactions. We also included a binary grouping variable in the temporal correlation structure based on river level (Fig 5, Step 2) because concentrations of NOx can vary more considerably during high flows than more stable flow periods (e.g. [38]). We did not know *a priori* what the most suitable cut off would be for the level-based AR(1) structure and so we tested three options for each site: (i) less than the first quartile (Q1), (ii) less than the median (Q2), and (iii) less than the third quartile (Q3; Fig 5, Step 3). We then implemented the two-stage model-selection procedure (Fig 5, Steps 4a,b), using backwards stepwise regression and cross-validation for NOx in the same general way as for TSS, except that models were fit separately to each site and three AR(1) structures were tested for each site. This produced nine models (3 sites x 3 AR(1) structures), which we then refit using REML to calculate a *cvRMSE* for each. For each site, the model with the lowest *cvRMSE* was deemed the best model, reducing the nine models to three. Although each of these three remaining models had the greatest predictive ability for the relevant site, our aim was to develop a single model that could be applied across all sites. Therefore, we composited all of the covariates from the three ‘best’ models to create a final model for NOx (Fig 5, Step 5). We refit this final model using the data from each site separately, using cross-validation to generate the *cvRMSE*, *cvPC* and *cvR^2^* and evaluate the predictive ability (Fig 5, Step 6).

#### Prediction using data from in situ sensors

After identifying the final models for TSS and NOx, we fit those models using covariates based on the data from the *in situ* sensors (i.e. turbidity, conductivity and/or level, as per the final TSS or NOx model structure), and used them to make predictions and associated estimates of uncertainty of TSS and NOx, respectively (Fig 5, Step 7). There was limited overlap in the timespans of the laboratory and *in situ* sensor data (Figs 2–4), and of course no sensor-measured observations of TSS or NOx on which to forecast future concentrations. Therefore, we generated the predictions and associated estimates of uncertainty using an infinite-horizon forecast [39], which assumes that forecasting is being made well into the future so that any short-term temporal correlations in the data (captured by the AR(1) structure) are irrelevant.

We used a leave-one-out cross validation (LOOCV) procedure to assess whether the final models for TSS and NOx fit to the sensor-measured surrogate covariates could accurately and precisely predict the response (Fig 5, Step7). This took a single observation from the sensor-measured covariate(s) as the validation dataset and the laboratory data minus a single observation (time-matched with the sensor validation observation) as the training dataset, which was fit using the final TSS or NOx model. A prediction was then made using the validation dataset, and the procedure was repeated for each observation. The laboratory and sensor observation were rarely synoptic, and so we used only those sensor measurements collected within one hour of a laboratory measurement (for MR, PR and SC respectively: n = 11, 49 and 28 for TSS; n = 11, 23, and 30 for NOx) as validation observations. Given the limited size of these data subsets, the LOOCV procedure was a more suitable method than the 5-fold cross-validation procedure [36] we used on the much larger, complete datasets of laboratory observations. We then calculated the *cvRMSE*, *cvPC* and *cvR^2^* using the ‘time-matched’ laboratory concentrations of TSS or NOx and the associated LOOCV predicted concentrations.

We performed all statistical analyses in R statistical software [40], using the nlme package [41] to implement the linear mixed-effects models. Turbidity, conductivity, TSS, NOx and river level were all log_10_-transformed prior to analyses to meet model assumptions. We chose the log_10_-transform because it is commonly used and has the benefit of being easy to interpret on the transformed scale. Predictions from the models were back-transformed with bias correction [42] for graphical visualization and assessment of accuracy and precision with respect to the laboratory TSS and NOx concentrations.

## Results

### TSS model based on laboratory data

The final TSS model explained over 90% of the variation in TSS (*cvR^2^*) and had excellent 95% prediction coverage (c*vPC* = 97.7%; Table 2; Fig 6) based on the 5-fold cross-validation. Although the linear relationship was strong, the model tended to under-predict at high TSS values, where data were relatively sparse (Fig 6). More specifically, at observed TSS concentrations greater than *c*. 100 mg/L, the model marginally over-predicted at MR, and under-predicted at both PR and SC (Fig 6).

**Table 2.**
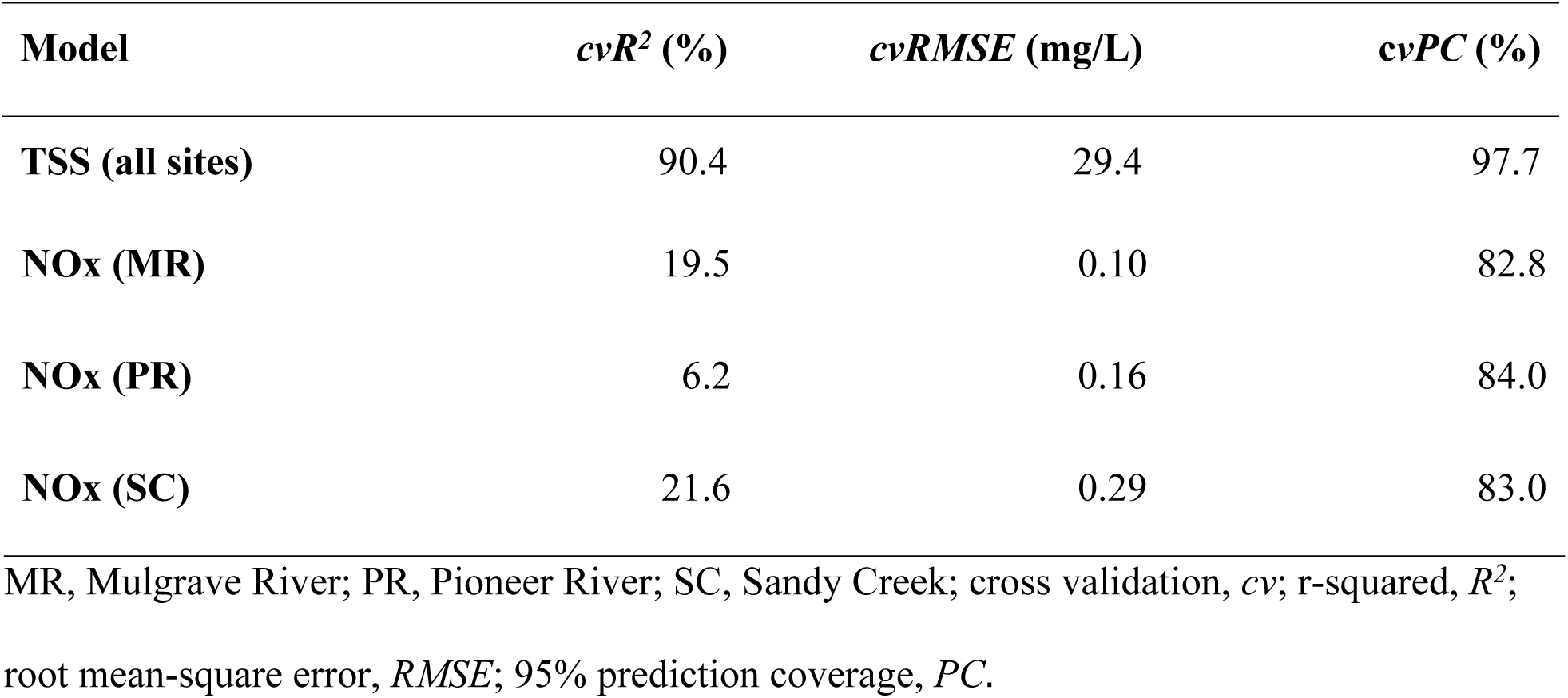
Cross validation statistics for the final total suspended solids (TSS) and final oxidized nitrogen (NOx) models as applied to data across all or each site.

| Model | *cvR*<sup>2</sup> (%) | *cvRMSE* (mg/L) | *cvPC* (%) |
| --- | --- | --- | --- |
| TSS (all sites) | 90.4 | 29.4 | 97.7 |
| **NO<sub>x</sub> (MR)** | 19.5 | 0.10 | 82.8 |
| **NO<sub>x</sub> (PR)** | 6.2 | 0.16 | 84.0 |
| **NO<sub>x</sub> (SC)** | 21.6 | 0.29 | 83.0 |

MR, Mulgrave River; PR, Pioneer River; SC, Sandy Creek; cross validation, *cv*; r-squared, $R^2$ ; root mean-square error, *RMSE*; 95% prediction coverage, *PC*.

**Fig 6.**
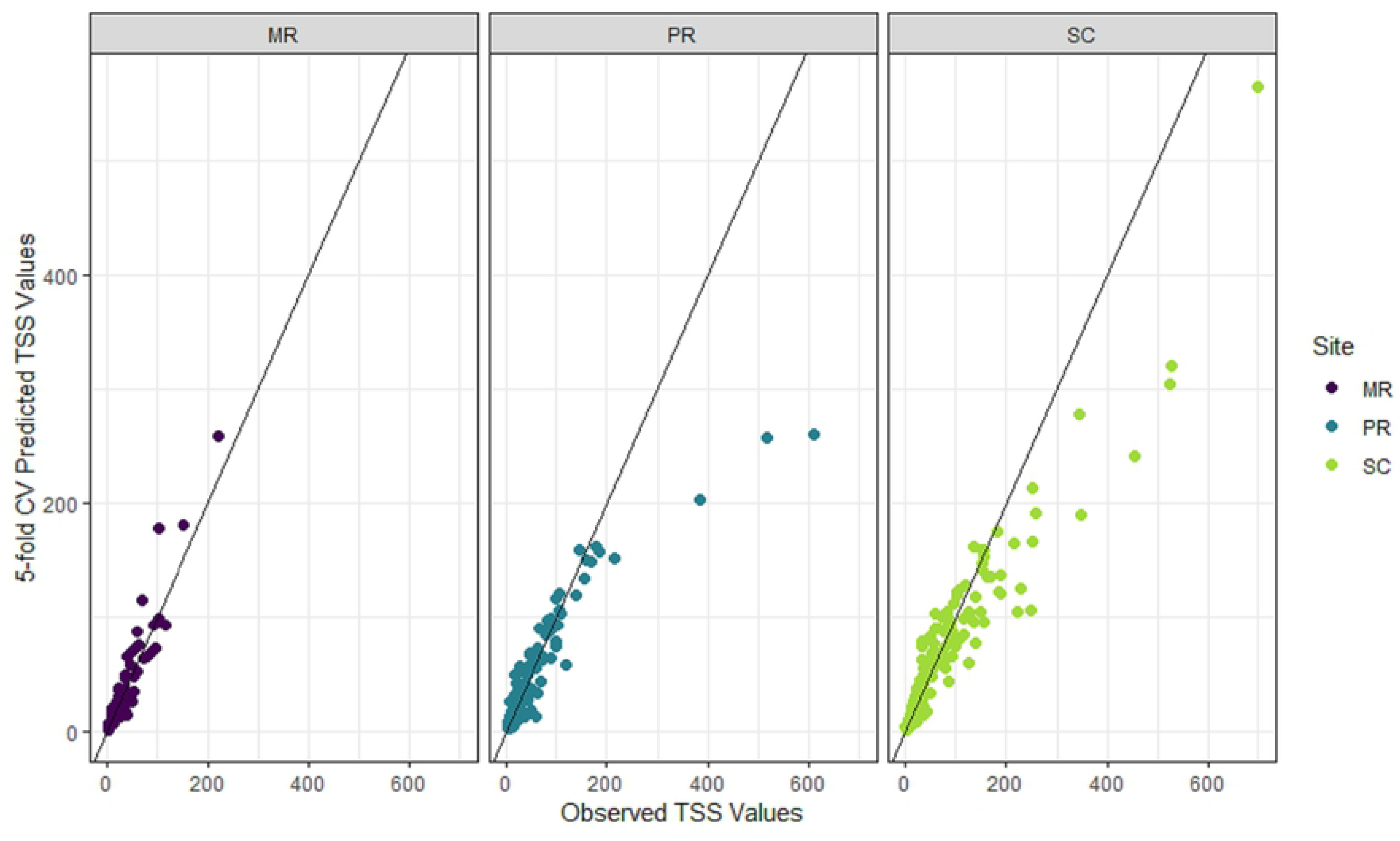
Observed versus 5-fold cross-validated (cv) prediction values of total suspended solids (TSS) from the final TSS model. TSS (mg/L; back-transformed with bias correction). Data from each site shown in purple (Mulgrave River, MR), blue (Pioneer River, PR) and light green (Sandy Creek, SC). Black lines show the 1:1 relationships between observations and predictions.

The relatively high correlation parameter value (*ϕ* = 0.87, with 95% confidence interval of 0.83-0.91) indicated that there was significant temporal autocorrelation in the data captured by the AR(1) model. The model included laboratory-based covariates for turbidity, site and T15, as well as interactions between site and turbidity and between T15 and turbidity (Table 3). TSS had a statistically significant (*p* < 0.05) and positive relationship with turbidity across all three sites, with TSS increasing more rapidly per unit rise in turbidity at MR than at the freshwater sites PR and SC, and when turbidity was ≥ 15 NTU as opposed to < 15 NTU.

**Table 3.** Coefficient estimates for the final TSS model fit to fit to laboratory data from all three sites.

| **Coefficient** | **Estimate** | **SE** | ***t*-value** | ***p*-value** |
| --- | --- | --- | --- | --- |
| **Intercept** | -0.28 | 0.098 | -2.86 | 0.0044 |
| **Turbidity** | 1.24 | 0.068 | 18.15 | < 0.0001 |
| **Site(PR)** | 0.11 | 0.069 | 1.59 | 0.1134 |
| **Site(SC)** | 0.002 | 0.082 | 0.024 | 0.9807 |
| **T15(Below)** | 0.41 | 0.089 | 4.60 | < 0.0001 |
| **Turbidity × Site(PR)** | -0.21 | 0.055 | -3.80 | 0.0002 |
| **Turbidity × Site(SC)** | -0.13 | 0.061 | -2.11 | 0.0354 |
| **Turbidity × T15(Below)** | -0.21 | 0.071 | -2.95 | 0.0034 |

TSS (total suspended solids, mg/L) and turbidity (NTU) were log<sub>10</sub>-transformed. MR, Mulgrave River; PR, Pioneer River; SC, Sandy Creek; SE, standard error.

### NOx model based on laboratory data

NOx models with a grouping structure based on the median value of river level (i.e. Q2) had the best predictive ability at all of the sites according to the *cvRMSE* (S1 Table). The combination of covariates from those models comprised turbidity, conductivity, level, and interactions between turbidity and level and between conductivity and level, which were all included in the final NOx model (Fig 7) along with median river level (Q2, as relevant to each site) in the grouping structure. Correlation parameters for this model indicated that temporal autocorrelation in the data was captured by the AR(1) structure (MR: *ϕ*= 0.86, with a confidence interval (CI) of 0.75-0.93; PR: *ϕ*= 0.86, CI = 0.72-0.94; SC: *ϕ*= 0.87, CI = 0.73-0.94).

**Fig 7.**
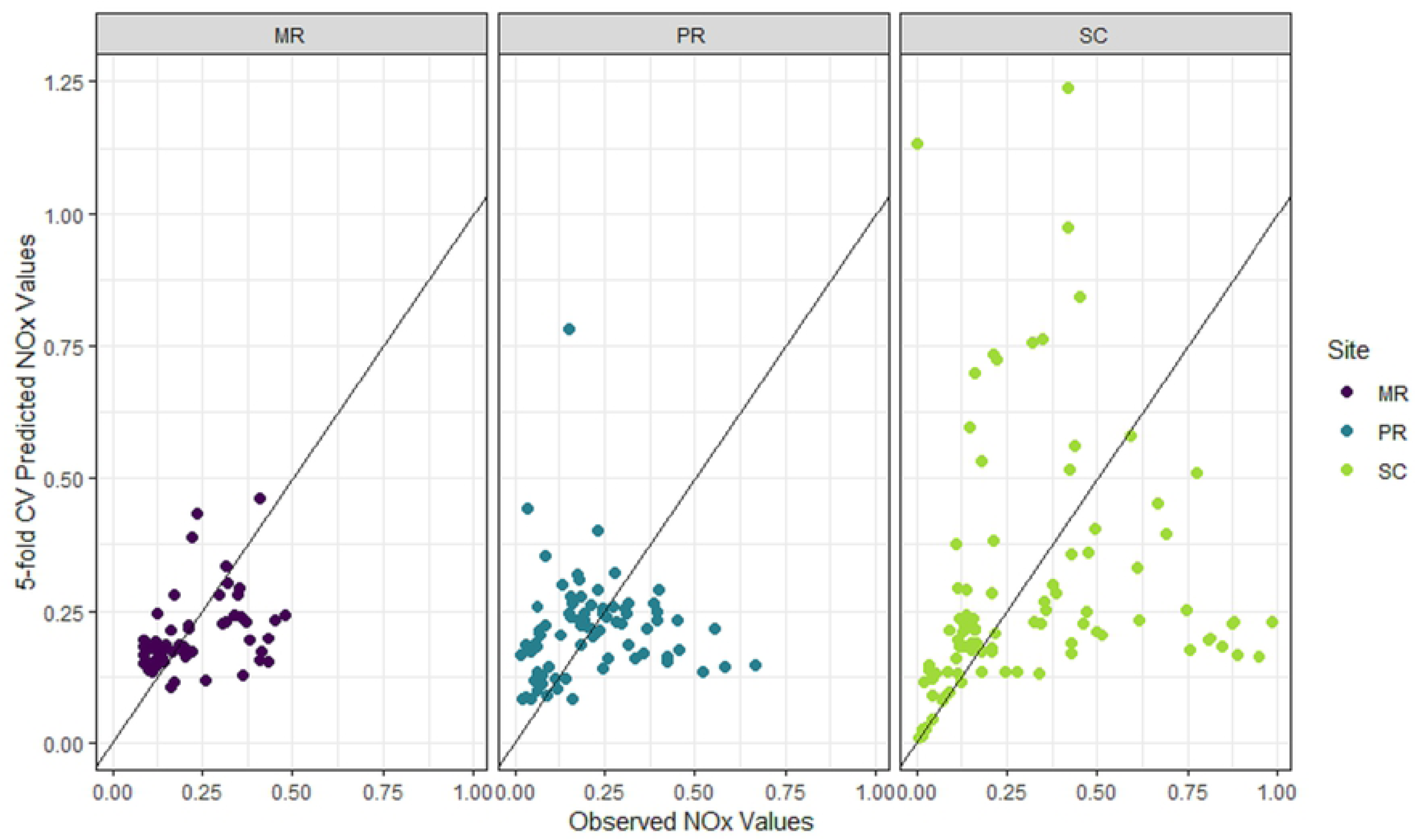
Observed versus 5-fold cross-validated (CV) prediction values of oxidized nitrogen (NOx) from the final NOx model. NOx (mg/L; back-transformed with bias correction). Data from each site shown in purple (Mulgrave River, MR), blue (Pioneer River, PR) and light green (Sandy Creek, SC). Black lines show the 1:1 relationships between observations and predictions.

The three site-specific models had marginal predictive ability, explaining 6 - 22% of the variation in NOx only (Table 2). At observed NOx concentrations greater than *c*. 0.1 mg/L NOx, the predictions had almost no relationship with the observations at any of the sites (Fig. 7). In addition, the statistical significance and direction of the covariates’ effects in the model also differed among sites (Table 4). For example, the relationship between conductivity and level was significant (*p* < 0.01) and positive at SC, but non-significant at MR and PR. The relationship between turbidity and level was significant (*p* < 0.01) and negative at PR, but non-significant at MR and SC. However, such differences may be expected given that the final model contained covariates and interactions were not significant at every site (S1 Table).

**Table 4.** Coefficient estimates for the final NOx model fit to fit to laboratory data from each site.

| Site | Coefficient | Estimate | SE | *t*-value | *p*-value |
| --- | --- | --- | --- | --- | --- |
| **MR** | Intercept | -24.22 | 16.35 | -1.48 | 0.1446 |
|  | Turbidity | -0.50 | 3.27 | -0.15 | 0.8799 |
|  | Conductivity | 14.99 | 9.12 | 1.64 | 0.1062 |
|  | Level | 20.92 | 15.93 | 1.31 | 0.1948 |
|  | Conductivity × Level | -13.47 | 8.87 | -1.52 | 0.1348 |
|  | Turbidity × Level | 0.50 | 3.21 | 0.16 | 0.8759 |
| **PR** | Intercept | -36.84 | 53.52 | -0.69 | 0.4933 |
|  | Turbidity | 18.88 | 6.00 | 3.15 | 0.0024 |
|  | Conductivity | 0.71 | 22.02 | 0.03 | 0.9745 |
|  | Level | 31.32 | 45.86 | 0.68 | 0.4967 |
|  | Conductivity × Level | -0.79 | 18.87 | -0.04 | 0.9668 |
|  | Turbidity × Level | -16.15 | 5.14 | -3.14 | 0.0024 |
| SC | Intercept | -2.42 | 0.69 | -3.50 | 0.0007 |
|  | Turbidity | 0.13 | 0.15 | 0.87 | 0.3884 |
|  | Conductivity | 0.64 | 0.24 | 2.68 | 0.0089 |
|  | Level | -4.37 | 1.41 | -3.10 | 0.0026 |
|  | Conductivity × Level | 1.47 | 0.47 | 3.09 | 0.0027 |
|  | Turbidity × Level | 0.62 | 0.33 | 1.84 | 0.0688 |

TSS (total suspended solids, mg/L), turbidity (NTU), conductivity ( $\mu\text{S}/\text{cm}$ ) and river level (m) were $\log_{10}$ -transformed. MR, Mulgrave River; PR, Pioneer River; SC, Sandy Creek; SE, standard error.

### TSS predictions from sensor data

The predictive accuracy of the final TSS model that included covariates from in situ sensors was high. The *cvR^2^* and *cvRMSE* (86.5% and 25.1 mg/L; Table 5, Fig 8) from the LOOCV showed that the accuracy of the model was similar to that of the model fit to laboratory data (*cvR^2^* = 90.4% and *cvRMSE* = 29.4 mg/L; Table 2). Although the 95% prediction coverage decreased from 97.7% to 88.6%, this is still reasonable given the relatively small sample size. Furthermore, all TSS predictions made from the sensor-measured turbidity covariate fell within the ranges of TSS measured in the laboratory (Table 1), except for some of the ‘future’ predictions at MR (Fig 9) when sensor-measured turbidity in late 2017 and early 2018 was high relative to that measured in the laboratory between January 2016 and June 2017 (Fig 2). The higher TSS values predicted at MR during the latter half of 2017 are therefore reasonable. TSS at SC was under-predicted at higher concentrations (> *c*. 100 mg/L; Fig 8), which was similar to our findings from the final TSS model fit to laboratory data (Fig 6). Finally, prediction intervals for all sites tended to be wider during peak events than at other times, which is to be expected when dealing with a log-normal response (Fig 9).

**Table 5.** Leave-one-out cross validation statistics for the final total suspended solids (TSS) and final oxidized nitrogen (NOx) models.

| Model | $cvR^2$ (%) | $cvRMSE$ (mg/L) | $cvPC$ (%) |
| --- | --- | --- | --- |
| **TSS (all sites)** | 86.5 | 25.1 | 88.6 |
| **NO<sub>x</sub> (MR)** | 56.9 | 0.05 | 100 |
| **NO<sub>x</sub> (PR)** | 6.6 | 0.10 | 100 |
| **NO<sub>x</sub> (SC)** | 71.1 | 0.11 | 100 |

Sensor-measured covariates were used as the validation dataset. Models applied across data from all (TSS) or each site (NO<sub>x</sub>). Cross validation, *cv*; r-squared, $R^2$ ; root mean-square error, *RMSE*; 95% prediction coverage, *PC*.

**Fig 8.**
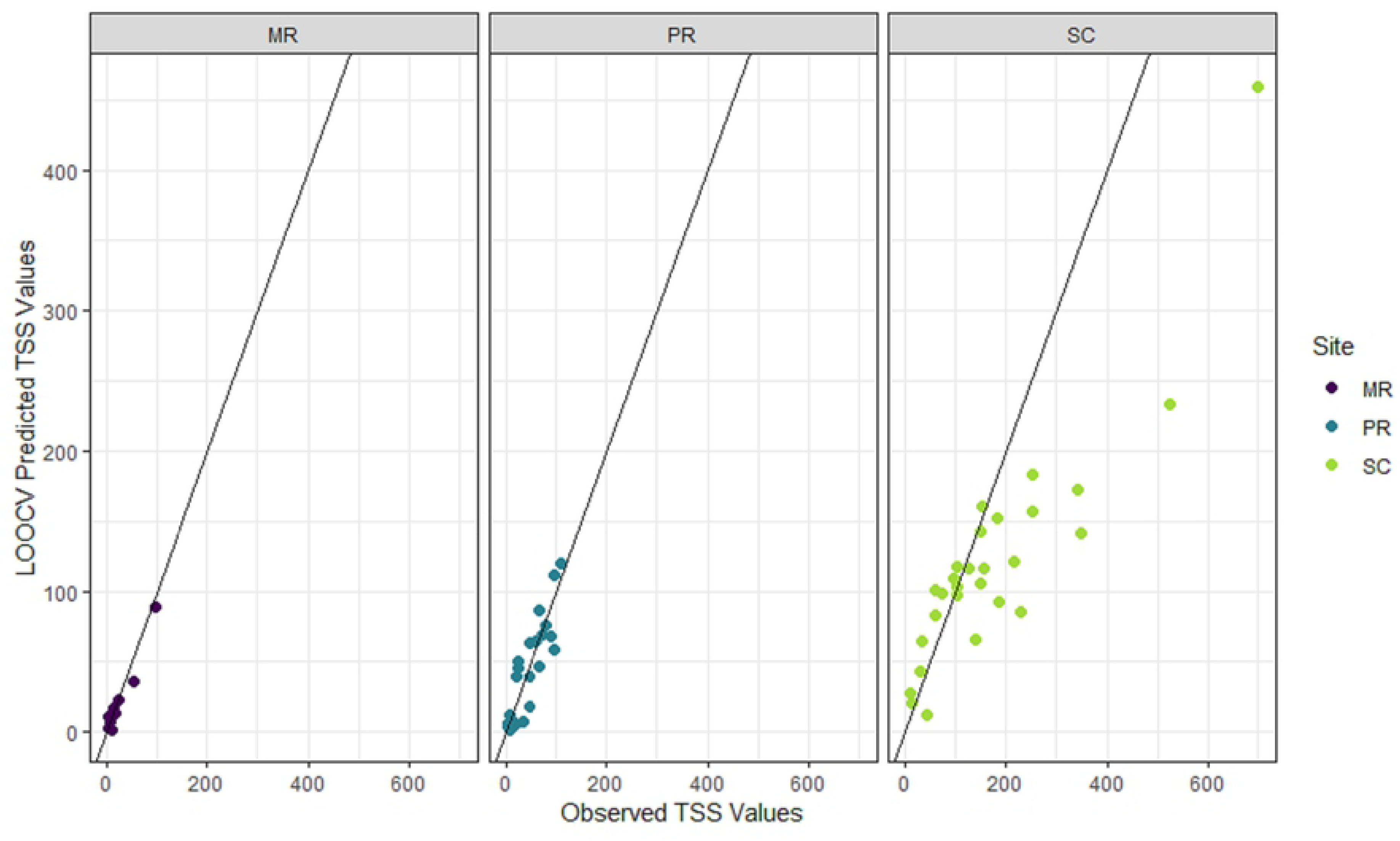
Observed total suspended solids (TSS, mg/L) versus predicted TSS (mg/L) from the leave-one-out cross validation (LOOCV).

**Fig 9.**
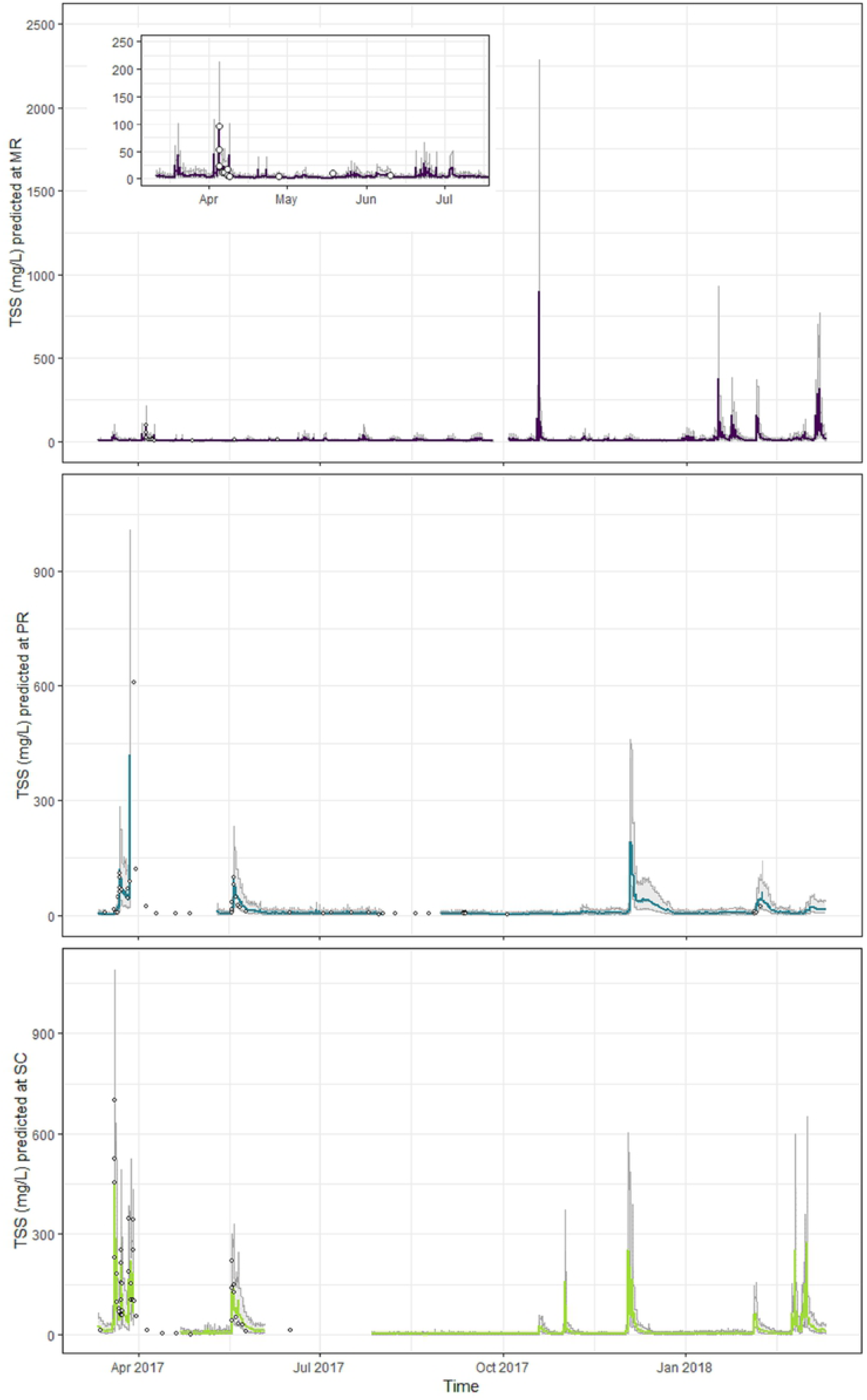
Total suspended solids (TSS, mg/L) at each site predicted using the final TSS model and *in situ* sensor turbidity data (March 2017-2018). Mulgrave River (MR, purple), Pioneer River (PR, blue) and Sandy Creek (SC, light green). Gray shading shows upper and lower boundaries of the 95% prediction interval, and the inner lines the predicted TSS concentrations through time. Gaps indicate periods of missing data in the sensor time series. Closed circles show the laboratory-measured TSS concentrations within the same period.

### NOx predictions from sensor data

Accuracy and precision of the NOx predictions produced by the model fit to the sensor-based covariates exceeded that of the final model fitted to all of the laboratory data (Tables 2 and 5, Fig 10). The *cvRMSE* values were substantially smaller (MR: 0.05 vs 0.10; PR: 0.10 vs 0.16; SC: 0.11 vs 0.29) and the *cvR^2^* values higher (MR: 56.9 vs 19.5%; PR: 6.6 vs 6.2%; SC: 71.1 vs 21.6%). In addition, the 95% prediction intervals were more reliable in the model fit to sensor data and captured the true values 100% of the time, with prediction intervals tending to be wider during events than non-events (Fig 11). At SC, there was a close relationship between the predictions and the observations at concentrations < *c*. 0.15 mg/L, but values greater than those tended to be under-predicted (Fig 10). There was also consistent over-prediction at MR and no relationship between predicted and observed NOx at PR (Fig 10).

**Fig 10.**
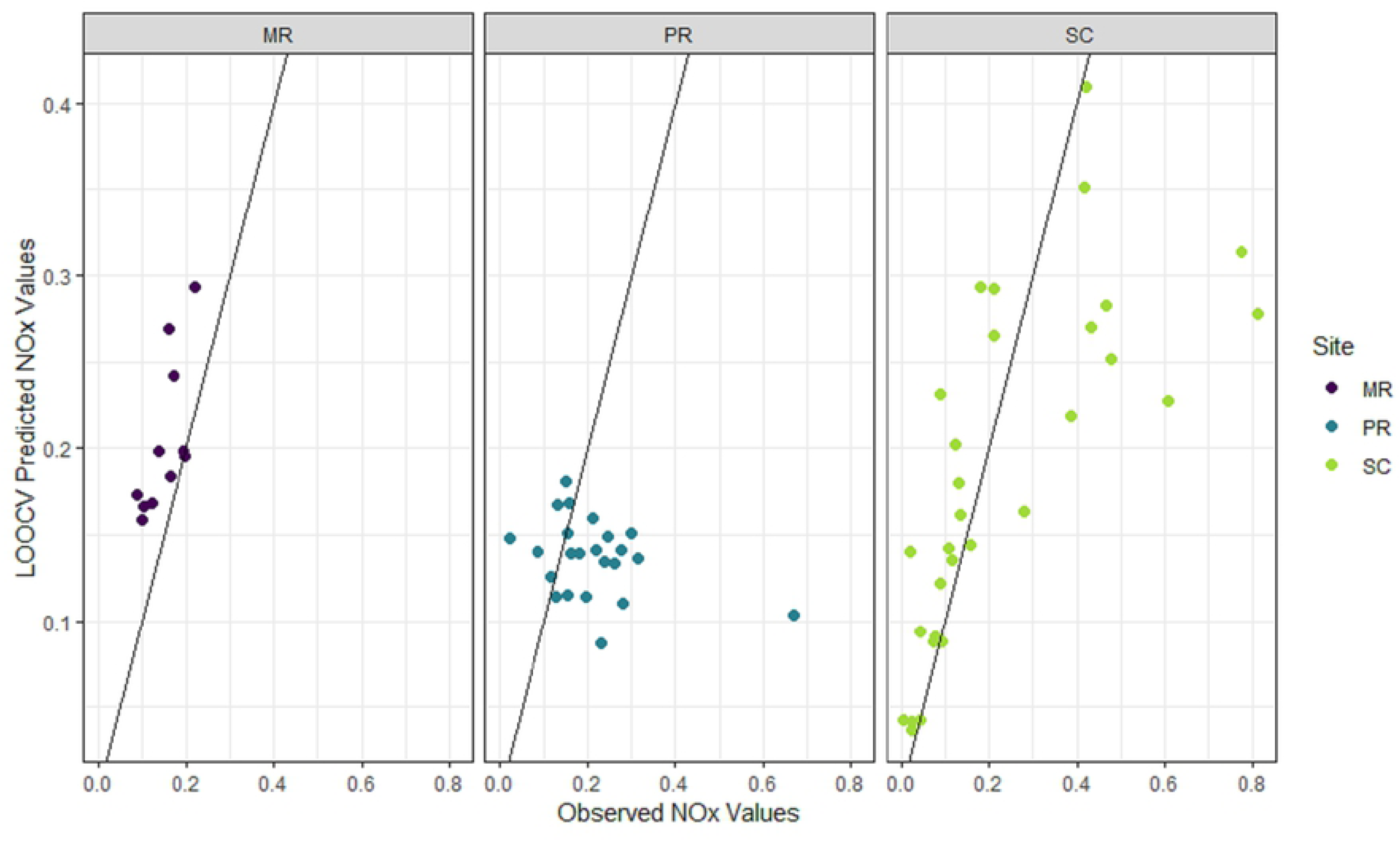
Observed oxidized nitrogen (NOx; mg/L) versus predicted NOx (mg/L) from the leave-one-out cross validation.

**Fig 11.**
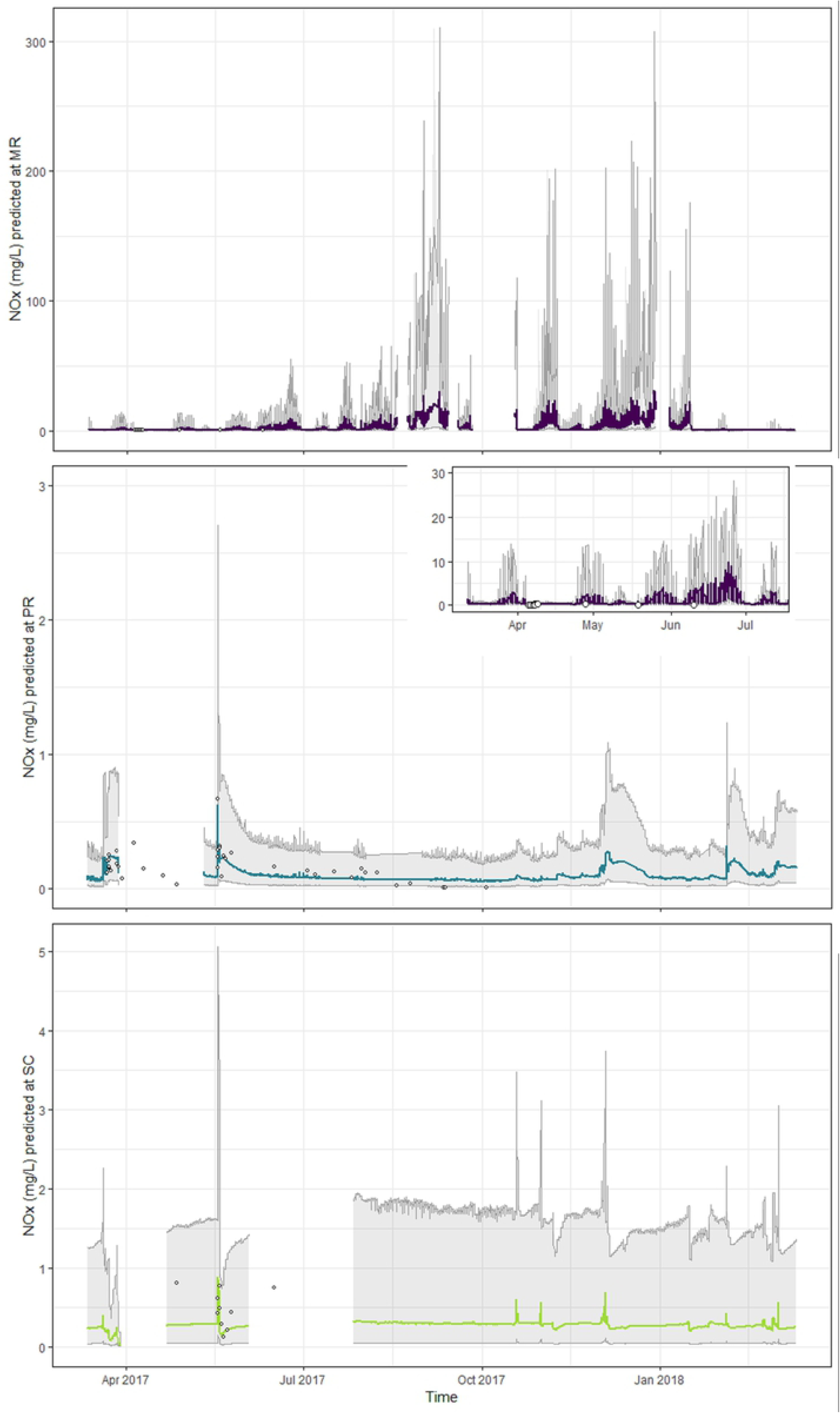
Oxidized nitrogen (NOx, mg/L; log_10_transformed) at each site predicted using the final NOx model and *in situ* sensor turbidity, conductivity and level data (March 2017-2018). Mulgrave River (MR, purple), Pioneer River (PR, blue) and Sandy Creek (SC, light green). Gray shading shows upper and lower boundaries of the 95% prediction interval, and the inner lines the predicted TSS concentrations through time. Gaps indicate periods of missing data in the sensor time series. Closed circles show the laboratory-measured NOx concentrations within the same period.

The range of NOx values predicted from the sensor covariates using all of the sensor-measured observations at both PR and SC fell within the range of NOx values measured in the laboratory. Although the maximum value predicted at MR (30.6 mg/L) was much greater than that measured in the laboratory (0.48 mg/L; Fig 11, Table 1), the elevated NOx values (>10 mg/L) predicted at MR were associated with peak values of the sensor-based covariates in late 2017 and early 2018 (turbidity, conductivity and/or level, depending on the site; Figs 2–4). As with TSS predictions, these high-concentration NOx predictions were in the ‘future’ (i.e. occurring after the last available laboratory observation); thus, the future NOx concentrations could conceivably have exceeded the concentrations observed in 2016 and early 2017.

## Discussion

### Surrogate potential and model generalizability

Here we predicted TSS and NOx from high-frequency water-quality data using models that accounted explicitly for temporal autocorrelation and heteroscedasticity, and that were developed specifically for their potential generalizability across the study sites. The transferability of models across sites, and potentially regions, will become increasingly important as organizations move to automated sensing for water-quality monitoring throughout catchments. We found a consistent, positive relationship between TSS and turbidity in both estuarine and fresh waters across study locations in different catchments separated by up to *c*. 700 km. In addition, the final TSS model had high predictive ability across all sites, indicating that a single mixed-effects model could be used to predict sediment concentrations from high-frequency, *in situ* sensor data over multiple locations in the study region. Whilst other studies have shown that turbidity is a useful surrogate of sediments in rivers, particularly when models account for temporal correlation in the data (e.g. [17]), our findings additionally suggest that a model based on sensor-measured turbidity has strong potential to be generalizable across locations, at least for the studied Great Barrier Reef catchments.

The complex relationships between NOx and its potential surrogates made the development of a generalizable model more difficult than it was for TSS. The predictive ability of the composite final NOx model fit to the turbidity, conductivity, and river level covariates at each site was substantially lower than that of the TSS model. Mismatch in the timing of laboratory and sensor observations was also a potential source of variability and bias in the leave-one-out cross-validations for the NOx model, particularly during events when rapid changes in the covariates’ values can occur. However, such mismatch would also have affected the leave-one-out cross-validations for the TSS model, which maintained good predictive ability regardless. Furthermore, the NOx model performed differently depending on the site to which it was applied, with a poor relationship between the observed and predicted concentrations at PR in particular, and a tendency to under-predict at SC for NOx concentrations above *c*. 0.1 mg/L. Consequently, we would not recommend that the NOx model developed herein be used as a generalized model across the study region.

The lack of generalizability of the NOx model likely relates to the complexity of dissolved nutrient dynamics in rivers, which are influenced by multiple and interacting factors including physical, chemical and biological processes [32,33]. For example, different timings and applications of fertilizers to agricultural land, different spatial configurations and types of soil and agricultural land (e.g. livestock grazing versus sugarcane cropland), and variation in the uptake of nutrients by phytoplankton, may all differentially influence dissolved nutrient concentrations among sites and through time [43,44,45]. Inclusion of additional covariates such as seasonal variation in fertilizer application, flow-weighted land use [46], soil characteristics and/or time since the last rainfall event may improve model fits and resultant predictions if more site-specific NOx models are desired. The NOx model findings also highlight that development of reliable, low-cost nitrate sensors [6] will remain an important management goal for the study region, and likely elsewhere, in the absence of suitable surrogate measures that can predict NOx across multiple sites with high accuracy and precision.

Important information for managers and scientists are also provided by the estimates of uncertainty associated with each prediction. Our results showed that the prediction intervals reliably captured the true TSS and NOx values for the final models fit to *in situ* sensor data, regardless of the predictive accuracy of the models. Prediction intervals were wider during events when the predicted TSS and NOx concentrations increased rapidly, corresponding with sudden new inputs of fresh water, and this has modelling implications. For instance, if such models were used to predict sediment and nutrient concentrations during events in the study region, end-users would need to be aware that the uncertainty around those predictions may be quite high, especially at the upper end of the prediction interval. Furthermore, if the predicted concentrations were then used to estimate high-frequency sediment and nutrient loads, as most water-quality monitoring programs transitioning to automated *in situ* sensors would desire, the associated estimates of uncertainty could be propagated through the model and accounted in loads estimates [47]. Knowledge of the magnitude of prediction uncertainty is important because it provides managers with information about where and when they can be most or least confident in model predictions [48] in order to prioritize future sampling efforts and management actions effectively [49,50]. We therefore recommend that measures of uncertainty (e.g. prediction intervals) be included routinely in water-quality reporting.

### Future directions and concluding remarks

Significant investments are being made to change management practices and reduce the quantities of sediments and nutrients entering rivers and, eventually, the Great Barrier Reef lagoon [25]. However, measuring the downstream impacts of these investments is challenging because current water-quality monitoring relies on a relatively small number of sites at or near river mouths. This makes pinpointing where the largest sources of sediments and nutrients are within a catchment difficult. The ability to predict TSS and NOx using data from relatively low-cost *in situ* sensors will allow networks of sensors to be deployed throughout catchments as technologies advance, creating numerous benefits for management. Firstly, the number of water-quality monitoring sites would increase significantly. Secondly, as the amount of data increases, the opportunity to develop near-real time statistical models for TSS and NOx increases, which could then be used to create dynamic predictive maps of sediment and nutrient concentrations throughout entire catchments. This would provide managers with greater situational awareness of where and when water-quality targets are being breached and would allow prioritization of land management actions in space and time to further reduce land-based impacts on the Great Barrier Reef lagoon.

Our study highlights that models fit to *in situ* water-quality data can be used to generate accurate predictions of TSS at both freshwater and estuarine sites. As the number of monitoring locations increases, spatial statistical models for stream networks [51] could be used to generate predictions, with estimates of uncertainty, across entire catchments. These methods could also be extended into both space and time (i.e. spatio-temporal models [52]), the need for which still clearly exists [31,53]. Such efforts, in combination with the methods developed herein, could revolutionize the way water quality is monitored and managed.

## Acknowledgments

The authors acknowledge the Queensland Department of Environment and Science, Great Barrier Reef Catchment Loads Monitoring Program for the data and the staff from Water Quality and Investigations for their input. We thank Grace Heron for producing the map in Fig 1. Laboratory and *in situ* sensor data together with the R code used in this paper are provided in the Supporting Information.

## Supporting information captions

**S1 Data and code.** Water-quality data files used to fit and predict from the TSS and NOx models using the R script herein.

**S1 Table. Site-based NOx models.** The best NOx model for each site based on the *cvRMSE* (i.e. lowest value per site), following backwards stepwise selection of the covariates in each model using the Akaike Information Criterion.

**S1 Figure. Turbidity and TSS.** The relationship between laboratory-determined turbidity (NTU) and total suspended solids (TSS, mg/L), log_10_-transformed, at Mulgrave River (MR; left plot), Pioneer River (PR, middle plot) and Sandy Creek (SC, right plot), with turbidity values < 15 NTU represented by closed triangles and those ≥ 15 NTU by closed circles, with each set in the freshwater sites PR and SC enclosed by ellipses.

**S2 Figure. Turbidity and NOx.** The relationship between laboratory-determined turbidity (NTU) and oxidized nitrogen (NOx, mg/L), log_10_-transformed, at Mulgrave River (MR; left plot), Pioneer River (PR, middle plot) and Sandy Creek (SC, right plot).

**S3 Figure. Conductivity and NOx.** The relationship between laboratory-determined conductivity (µS/cm) and oxidized nitrogen (NOx, mg/L), log_10_-transformed, at Mulgrave River (MR; left plot), Pioneer River (PR, middle plot) and Sandy Creek (SC, right plot).

**S4 Figure. River level and NOx.** The relationship between river level (m) recorded on-site at the time of water sample collection and oxidized nitrogen (NOx, mg/L), log_10_-transformed, at Mulgrave River (MR; left plot), Pioneer River (PR, middle plot) and Sandy Creek (SC, right plot).

**S5 Figure. TSS-model residuals.** Residuals from the final total suspended solids (TSS, mg/L, log_10_-transformed) model plotted in chronological order of the observations at Mulgrave River (MR; left plot), Pioneer River (PR, middle plot) and Sandy Creek (SC, right plot).

**S6 Figure. TSS-model diagnostic plots.** Diagnostic plots for the final total suspended solids (TSS, mg/L, log_10_-transformed) model. Upper row: fitted values vs residuals. Middle row: boxplots of residuals. Lower row: QQ-plot. Left column: by site (MR, Mulgrave River; PR, Pioneer River; SC, Sandy Creek). Right column: by T15 (Above: turbidity ≥ 15 NTU; Below: turbidity < 15 NTU).

**S7 Figure. NOx-model residuals.** Residuals from the final oxidized nitrogen (NOx, mg/L, log_10_-transformed) models plotted in chronological order of the observations at Mulgrave River (MR; upper plot), Pioneer River (PR, middle plot) and Sandy Creek (SC, lower plot).

**S8 Figure. NOx-model diagnostic plots.** Diagnostic plots for the final oxidized nitrogen (NOx, mg/L, log_10_-transformed) model for each site (Mulgrave River, MR; purple points; Pioneer River, PR; blue points; Sandy Creek, SC; light green points). Upper row: fitted values versus residuals. Middle row: boxplots of residuals. Lower row: QQ-plots.

